# Simplest Model of Nervous System. I. Formalism

**DOI:** 10.1101/2023.11.23.568481

**Authors:** Anton V. Sinitskiy

## Abstract

This study presents a novel, highly simplified model of the nervous system, inspired by one hypothetical scenario of its origin. The model is designed to accommodate both mathematical derivations and numerical simulations, offering a template for studying generalized principles and dynamics beyond the specifics of the referenced origin scenario. The model offers a holistic perspective by treating the nervous system and the environment (in their simplest forms) as parts of one system and, together with a companion paper, notes the key role of evolutionary factors (in this model, predator evasion) in shaping the properties of the nervous system. To emphasize these fundamental principles, some aspects, such as the highly dimensional nature of the networks or detailed molecular mechanisms of their functioning, are omitted in the current version. Analytically, the model facilitates insights into the stationary distribution as a solution to the Fokker-Planck equation and the corresponding effective potential and rotation (solenoidal) terms. Numerically, it generates biologically plausible (given its high abstraction) solutions and supports comprehensive sampling with limited computational resources. Noteworthy findings from the study include limitations of the commonly used weak noise approximation and the significance of rigorous mathematical analysis over heuristic interpretations of the potential. We hope that this abstract model will serve as a fruitful tool for better understanding a complete set of principles for modeling nervous systems.

## Introduction

“Everything should be made as simple as possible, but not simpler.” Even though Einstein might have never said it,^1^ we follow this principle for our study. We aim to construct a simple (perhaps the simplest) model for the nervous system that captures, without oversimplifying, its essence.

Models which distill intricate phenomena into simpler forms have historically played a pivotal role in various scientific disciplines. For instance, the model of a hydrogen atom in Quantum Chemistry not only illustrates the fundamental principles of Quantum Mechanics in application to atoms, but also offers foundational concepts, such as orbitals, for the approximate modeling of more complex molecules. Similarly, the Lorenz model in meteorology captures key phenomena like chaotic behavior, even though it significantly simplifies the intricate details of its original meteorological counterpart. We believe that neural network studies can similarly benefit from such models. Though numerous models exist in Theoretical and Computational Neuroscience, our objective here is to provide a novel, comprehensive yet simple model to help formulate and test general principles of and specific approaches to studying neural networks. We also hope that the resulting insights might be partially extended to studying not only biological, but also artificial neural networks.

For context, we draw inspiration from a hypothetical scenario regarding the evolution of the nervous system, as recently proposed by Paulin and Cahill-Lane.^2^ They suggest that a primitive nervous system could arise in *Dickinsonia* or similar animals in the Ediacaran period (we will further refer to them as “Dickinsonia” in quotes). They fed on organic matter from bacterial mats through external digestion, occasionally moving from one area of the seabed to another (Fig. 1a). If one organism overlaid another, it could lead to significant damage to or even death of the latter due to external digestion. The enhancement of the sensitivity of some peripheral cells to electric fields created by other “Dickinsonias” and triggering an escape reaction of the entire “Dickinsonia” by such sensitive cells should have been supported by natural selection. In this way, a primitive nervous system might have evolved as a survival mechanism.^2^ It is crucial to underline that this scenario remains speculative; even the existence of a nervous system in *Dickinsonia*, or *Proarticulata* in general, remains unconfirmed. Existing reconstructions of the origin of the nervous system are incomplete, and no consensus exists on what organisms were the first to develop it.^3-11^ Yet, we believe that the mathematical model built upon such a premise will hold generalizable value.

**Fig. 1.**
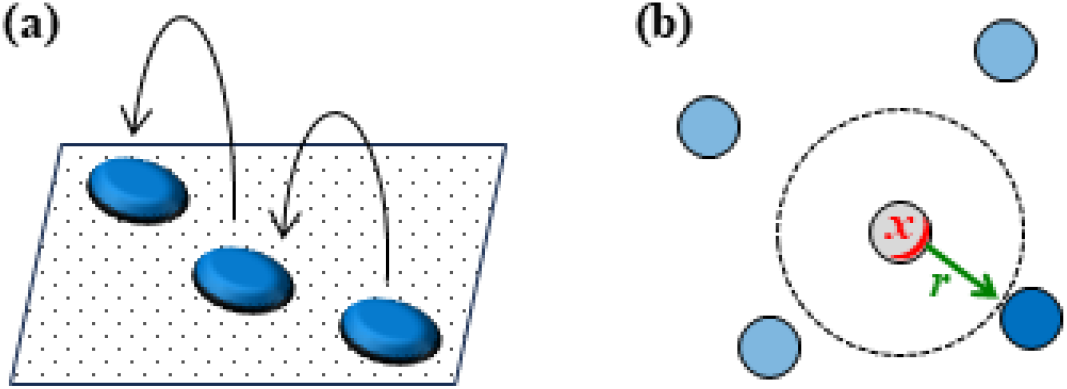
Motivating our mathematical model is a hypothetical scenario detailing the emergence of a primitive nervous system. (a) *Proarticulata* fed by external digestion of bacterial mats occasionally moving on the sea floor.^12^ Sporadic overlaying of one organism by another could cause significant damage, and the nervous system might have arisen as an adaptation to evade such events.^2^ (b) Our model includes only two variables: the distance *r* (*green*) between the organism (*grey, in the center*) and the nearest predator (*blue*), and a variable *x* (*red*) that describes the excitation of the nervous system, possibly interpreted as the membrane potential of the nearest neuron, or an average value of the potentials of all neurons in the network.

Reference to this specific hypothetical scenario offers a fresh perspective and allows us to go beyond the existing models. For example, current literature often presents the function of a biological neural network as something like an unsupervised learning of probability distributions of the external world to minimize sensory uncertainty or surprisal^13-20^ or, even simpler, as a minimization of mean square errors.^21-25^ But if we bear in mind the hypothetical scenario of “Dickinsonia”, then it is easier to notice that the connection between the sensory uncertainty or mean square error and evolutionary fitness is not straightforward and requires an explicit consideration. Similarly, while commendable strides have been made in constructing a general neural network theory,^21,22,26-30^ we are worried, after looking at them from the viewpoint of this hypothetical evolutionary scenario, that such efforts might be at risk of leading to mathematical constructs applicable to anything in general, but to nothing in particular. In this work, we chose the path of drawing from a specific evolutionary scenario to ensure that we do not overlook essential components that may be easy to miss in a general picture.

Our model^31^ aims to address several key questions. Is it feasible to apply recent formalisms from nonequilibrium statistical physics to the dynamics of a nervous system in a specific environment? If so, what potential function and rotation (solenoidal) terms manifest in the plausible dynamics of such a system? How does the neural network dynamics intertwine with, and get constrained by, the dynamics of its environment? In modeling, what are the implications of assuming infinitesimally weak noise, and how does this assumption align with the biological interpretation of our model? Furthermore, we extend the scope of the analysis to the questions of evolutionary optimization in a companion paper.^32^

## Methods

### Dynamical model

To describe the most significant aspects of the dynamics of the “Dickinsonia” and its environment (other “Dickinsonias”, considered in this case as predators), two dynamic variables suffice: *x*(*t*), which shows how the neuronal membrane potential changes over time *t*, and *r*(*t*), which expresses the distance to the nearest predator (Fig. 1b). Of course, one “Dickinsonia” could contain many neurons; in this simple model, we assume that *x* is the potential of the neuron closest to the predator, and therefore the most strongly reacting (alternatively, it might be interpreted as the aggregated activity of several neurons closest to the predator or of all neurons in the nervous system). Similarly, *r* is a one-dimensional simplified description of the complex spatial distribution of predators around a given “Dickinsonia”. Simplistically, *r* is the distance to the nearest predator.

Next, we assume that the rate of change of these variables over time depends only on their current values, and also includes random noise:

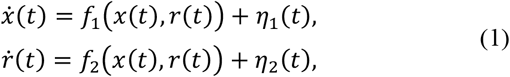

where 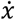 and 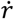 are the derivatives of *x* and *r* over time, *f*_1_ and *f*_2_ are some functions of *x* and *r*, and *η*_1_ and *η*_2_ are random variables (noise). As a biologically reasonable simple form for *f*_1_ and *f*_2_, the rate of change of *x* over time may include a term that returns the membrane potential *x* to the resting potential *x*_0_, and a term that expresses the neuron’s sensitivity to the predator’s approach:

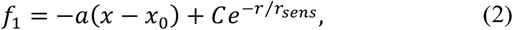

where *a, C* and *r*_sens_ are positive constants. The rate of change of *r* over time can include a term due to which exceeding the membrane potential *x* over the resting potential *x*_0_ triggers the organism’s escape from the predator, and a term that prevents the predator from moving infinitely far away (one predator can move very far, but *r* is the distance to the nearest predator; for the derivation of the corresponding effective potential and force, see Appendix A):

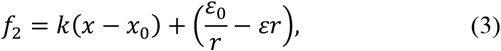

where *k, ε* and *ε*_0_ are positive constants. Finally, regarding the random variables *η*_1_ and *η*_2_, the standard assumptions seem reasonable that they obey Gaussian distributions with zero mean values (otherwise non-zero means could be included in the definition of *f*_1_ and *f*_2_) and the following covariances:^27,33-39^

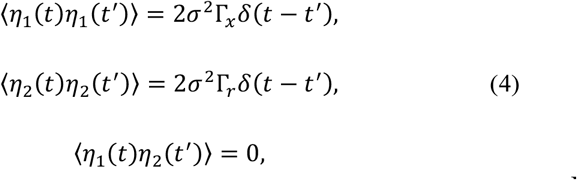

where *σ*^2^ sets the scale of random fluctuations and in the course of further mathematical analysis can be interpreted as a small parameter, common for heterogeneous variables *x* and *r* (in similar stochastic equations describing physical systems, *σ*^2^ is the temperature in units of energy), *δ*(·) is the Dirac delta function, and Γ_*x*_ and Γ_*r*_ are positive constants. In other words, we assume that random fluctuations in the rate of change of the membrane potential and random fluctuations in the rate of change of the distance to the nearest predator are caused by different reasons and occur independently of each other, and that the magnitudes of both types of fluctuations do not depend on the current values of the potential and distance.

Thus, the simplest model of the simplest nervous system, with one variable *x* characterizing the current state of the nervous system, and one variable *r* characterizing the external world and the organism’s position in it, is given by equations (1) and (4). Integration of dynamical equations (1) provides stochastic dynamical trajectories of the system *x*(*t*) and *r*(*t*), which, in their turn, can be used to compute various properties of interest.

We expect that the scope of this model is quite wide, much beyond the mentioned hypothetical scenario of the emergence of the nervous system,^2^ and beyond a specific choice of the functions *f*_1_ and *f*_2_, as given by equations (2) and (3).

### Steady state and parameterization of the dynamics

To formulate and solve the problem of the evolutionarily optimal simplest neural network, it is convenient to use the toolkit for the analysis of stochastic equations of the form (1), developed in the context of nonequilibrium statistical physics and also previously applied to the analysis of functioning neural networks (but not their evolutionary optimization).^18,27,28,34,35,37-44^

Let us move from considering an individual trajectory of one instance of the system to considering the probability distribution of trajectories of the ensemble of systems. Let *P*(*x,r,t*) be the probability density of such distribution, that is, the probability of finding the given system at time *t* with values of the first variable in the range from *x* to *x+dx* and with values of the second variable in the range from *r* to *r+dr* equals *P*(*x,r,t*)*dxdr*. As is well-known, the change in *P* over time is described by the Fokker-Planck equation:

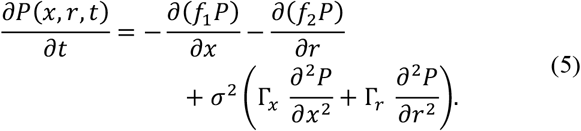

If the probability distribution over time reaches a stationary (unchanging with time) state, then the probability density in such a state *P*_*stat*_(*x,r*) is determined by the condition

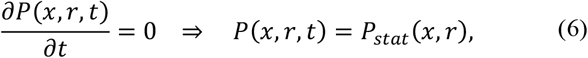

or, taking into account equation (5),

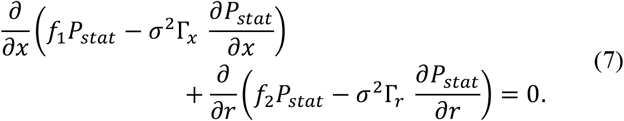

Next, we note that this equation can be rewritten, without loss of generality, as a system of two equations,

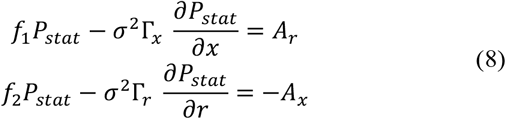

where *A* = *A*(*x,r*) is some function essentially defined by equations (8), and subscripts *x* or *r* here and below refer to partial derivatives with respect to the corresponding variable. In principle, equations (8) would suffice to derive all the results of this paper; however, for the purpose of agreement with the notations in the previous literature, we rewrite them in terms of a potential function *u*(*x,r*), which is an analogue of the common potential function in statistical physics generalized to nonequilibrium systems:

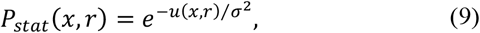

and a new function *Q* = *Q*(*x,r*) defined as:

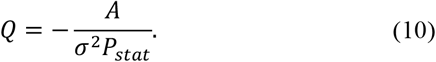

In these notations, equations (8) are rewritten as follows:

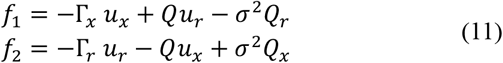

In the literature, it is common to take (sometimes implicitly) the limit of infinitely weak noise (*σ*^2^ → 0).^18,27-29,35,36,40,43-45^ In this case, equations (11) formally simplify to the following equations:

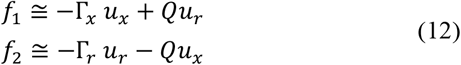

Known in the literature as the Helmholtz, Helmholtz-Ao or Helmholtz–Hodge decomposition.^27-29,35,40,42,43^ The question of the applicability of this approximation remains open (see Results).

Note that the function *Q* (or *A*) is a critically important component of a model of a nervous system, ensuring its nonequilibrium dissipative nature. Vanishing *Q* in physical applications corresponds to microscopic reversibility. In our model, if *Q* vanishes, then, taking into account equation (11),

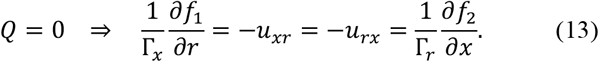

The partial derivative ∂*f*_1_/∂*r* shows the neuron’s responsiveness as a sensor to the distance to the predator. This partial derivative, by its biological interpretation, should be *negative* [see, for example, the second term in equation (2)]: the farther the predator, the weaker the sensory signal. On the other hand, the partial derivative ∂*f*_2_/∂*x* shows how the activation of the neuron affects the distance to the predator. Biologically, this derivative should be *positive* [see, for example, the first term in equation (3)]: neuron activation should signal to the effectors that they need to turn on, so that the organism moves away from the predator. These two requirements cannot be met simultaneously if *Q* = 0, as shown by equation (13). The biologically meaningful signs of these two partial derivatives are only possible in a nonequilibrium, dissipative system. In the model under consideration, this requirement means that *Q*(*x, r*) ≠ 0.

## Results

### Numerical simulations demonstrate the biological plausibility of the model

First of all, we estimated the relevance of the model by numerically integrating the stochastic differential equations (1) with functions *f*_1_ and *f*_2_ and random variable variances given by equations (2)-(4). Due to the high level of abstraction in the model and its motivation by a hypothetical evolutionary scenario, it would be difficult to set the values of the coefficients *a, C, k, ε*, Γ_*x*_ and Γ_*r*_ based on experimental or empirical data. (Note that the parameter *x*_0_ can be chosen arbitrarily, because the model equations are invariant to a constant translation in *x*. In numerical simulations, we used *x*_0_ = −70 mV, but we plotted the results relative to *x*_0_ to make this invariance explicit.) In the first round of simulations, we set all these parameters equal to 1. However, the generated trajectories were too noisy, and the expected pattern that the nervous system serves to evade predators did not clearly emerge. We then attempted minimal changes to the parameters, trying to find a regime where this behavior could be seen. This goal can be easily achieved, for example, with the following parameter values, which we used for the subsequent calculations:

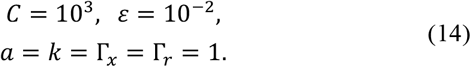

A typical trajectory [*r*(*t*), *x*(*t*)] generated with these parameters is shown in Fig. 2. The panel in Fig. 2a shows the central 1/10^th^ part of the trajectory fragment shown in Fig. 2b; the latter, in turn, shows the central 1/100^th^ part of the trajectory fragment from Fig. 2c. Finally, the trajectory fragment shown in Fig. 2c is the initial 1/100^th^ part of a full trajectory generated by numerical integration. This visualization aims at demonstrating the dynamics of the model on various timescales. The units for *r* (such as mm, etc.) and *x* (such as mV, etc.) depend on the choice of the units of the coefficients *a, C, k, ε*, Γ_*x*_ and Γ_*r*_, and we do not specify them here for the same reason: an indirect connection between the model and experimental data. Instead, we assume that *r* and *x* are expressed in some relative units, which does not affect the further analysis.

**Fig. 2.**
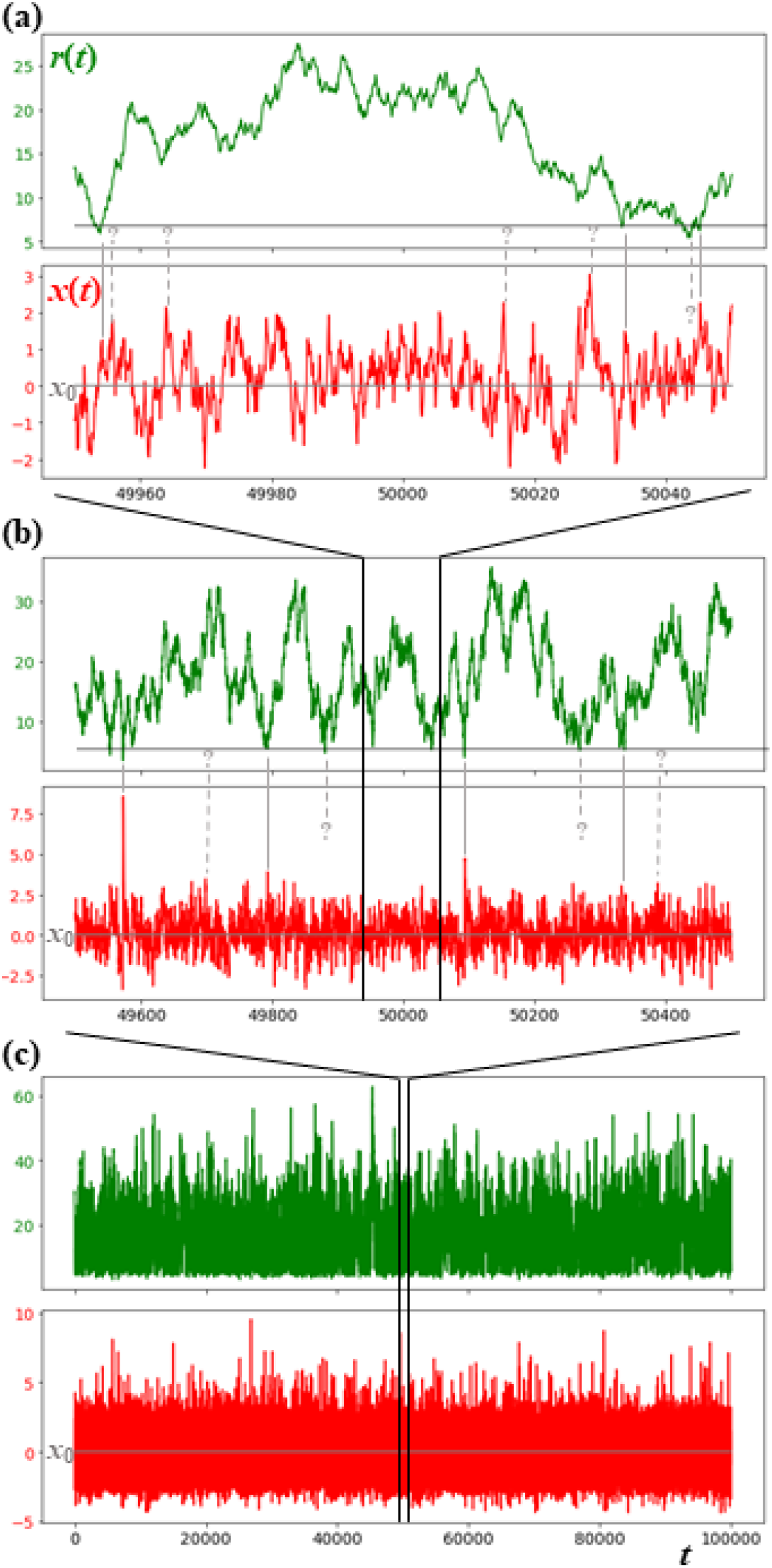
Numerical simulation demonstrates biological plausibility of the model. Dynamics of the distance *r* to the nearest predator (*green*) and the neuronal membrane potential *x* (*red*), found by numerical integration of equations (1)-(4) with the initial guess of the parameters (14), is biologically plausible, given the simplicity of the model. (a,b) On shorter timescales, local maxima of the membrane potential often coincide with local minima of the distance to the predator (*vertical grey lines*, biological interpretation: a nervous system excites in danger), and the distance to the predator after such events often increases (escape behavior); however, in other cases, such coincidence does not take place (*dashed lines with question marks*: unreliable functioning of a primitive nervous system). (c) On longer timescales, a stationary distribution of *x* and *r* is reached. The model nervous system is highly noisy, with recurrent events of neuronal excitation beyond the regular noise level. Such spike-resembling events seem instant on longer timescales (b,c), but reveal a finite width and certain structure at higher resolution (a).

On shorter timescales (∼10^2^-10^3^ units of time), the dynamics of both *x* and *r* is highly noisy (Fig. 2a,b). Despite this noise, often, but not always, the neuron tends to excite when a predator approaches, as seen from the comparison of the local minima of *r*(*t*) and the local maxima of *x*(*t*) (grey vertical lines in Fig. 2a,b). A connection is also observed between the neural excitation and the avoidance behavior (the work of effectors): after the strongest peaks in the neural activity, the distance *r* significantly increases.

On longer timescales (Fig. 2c, 10^5^ units of time), both *x* and *r* still look noisy, but stationary variables. The values of *r* fluctuate in the range of ∼5 to ∼50 relative units (typically, within the range of 10 to 25). As for the values of the membrane potential *x*, they typically lie in the range of ∼3-4 relative units above or below the resting potential *x*_0_. However, large positive deviations from *x*_0_ are evidently preferred over large negative deviations, which is biologically meaningful: depolarization occurs more frequently and on a larger scale than hyperpolarization in this model, due to the positive value of the parameter *C*.

Note that the plots for the membrane voltage *x* on longer timescales seem to have narrow peaks (Fig. 2b,c), corresponding to the scenario of sharp neuronal spikes, while on shorter timescales, they have a finite width and a certain shape (Fig. 2a), which more resembles a graded potential response in non-spiking neurons. As demonstrated in a companion paper,^32^ adjusting the numerical parameters in the model may make neuronal peaks look sharper (more ‘spiky’). Therefore, despite the simplicity of the model, it may express to some extent a spiking or non-spiking nature of a neuron.

Overall, the results presented in this subsection, in our opinion, may correspond to an early stage of evolution of a nervous system, namely preadaptation of “Dickinsonia” to predator avoidance, based on pre-existing excitable cells that somewhat reacted to the electric field of an approaching predator (and in this context started working as sensory neurons) and more often activated pre-existing cilia than deactivated them (respectively, becoming effectors). Before these structures were evolutionarily optimized, they perhaps had demonstrated unreliable, noisy functioning, evolutionarily advantageous only on average, over a large number of events. The noisy character of the trajectories generated by the model with the given values of parameters, revealing only a weak avoidance behavior, as well as the model’s ability to demonstrate such a behavior in a wide range of values of the parameters (data not shown), corresponds in our opinion to a preadaptive state of an early nervous system. From an initial state like this, evolution should have quickly improved the performance of early nervous systems. The question of evolutionary optimization in the presented model is addressed in a companion paper.^32^

### Stationary distribution exists

To verify the theoretical prediction on the existence of the stationary distribution, we ran multiple independent numerical simulations of the model, using different initializations of the random number generator. All generated individual trajectories [*x*(*t*),*r*(*t*)] were different from each other, but the corresponding probability distributions were the same (up to the numerical accuracy, which is limited by the finite temporal length of the trajectories). In particular, the probability density for *r*, regardless of the values of *x*, defined as

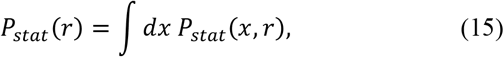

was found to be essentially the same from the two longest trajectories we ran (Fig. 3a, run 1: *green curve*, run 2: *green dots*). Both trajectories had the length of 10^7^ units of time, and the first 1/100^th^ of trajectory from run 1 is plotted in Fig. 2c. Similar distributions were obtained from other independent runs of shorter durations (data not shown).

**Fig. 3.**
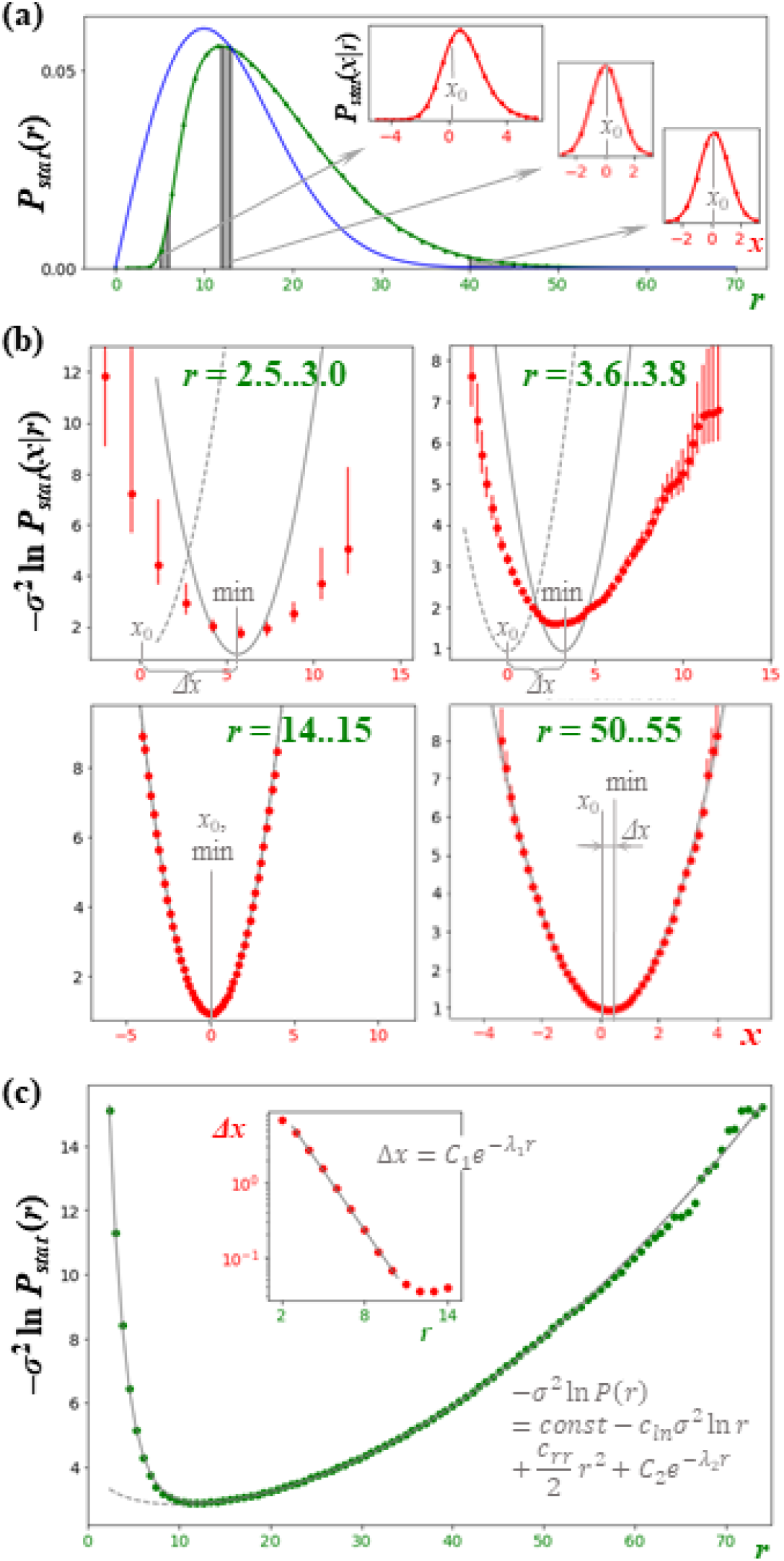
Stationary distribution *P*_*stat*_ exists and can be approximated analytically. (a) Distributions over *r* for two independent simulations are indistinguishable (run 1: *green curve*, run 2: *green dots*), but different from the distribution without the nervous system (*blue*) due to the active evasion of predators. Conditional distributions over *x* for different ranges of *r* are also indistinguishable for two independent simulations (inset, run 1: *red curve*, run 2: *red dots*). (b) Distributions over *x* can be approximated analytically (numerical results: *red, with 5% error bars*; long-distance asymptote (22): *grey dashed*; analytical approximation (30): *grey solid*). (c) Distributions over *r* can also be approximated (numerical results: *green dots*; analytical approximation (32): *grey solid*; the same, but without the exponential term: *grey dashed*). Inset: shifts in the minima of the distributions over *x* can be approximated by an exponential function of *r*. Note the negative log scales for the probability densities in (b,c) which are used to connect them to the components *u*_1_ and *u*_2_ of the potential *u*, respectively.

This probability density significantly differs from the probability density over *r* in the case of random distribution of predators in the environment, when a nervous system does not function [Fig. 3a, *blue curve*, plotted based on equation (48) from Appendix A]. The active avoidance of predators shifts the distribution toward larger values of *r*. In particular, the probability density in the region of small distances to the predator drops dramatically, while the tail of the distribution at large distances becomes wider. The most probable distance also shifts from 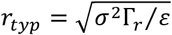 (see Appendix A), which equals 10 with the chosen values of the parameters, to a larger value, though this change is not as dramatic as the avoidance of small values of *r*.

The probability densities of *x* given the value of *r*, defined as

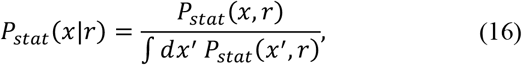

are also the same for independent runs of the simulations (Fig. 3a, insets, run 1: *red curves*, run 2: *red dots*; three insets correspond to *r* in the ranges of 5 to 6, 12 to 13 and 40 to 45, respectively).

These conditional probabilities of *x* follow Gaussian distributions at large or moderate distances to the predator *r*, while at short distances *r*, the distributions are wider and non-Gaussian, biased toward values *x* > *x*_0_ (Fig. 3a, insets). In particular, for run 1, with the given amount of sampling, the Kolmogorov-Smirnov test detects non-Gaussianity of *P*_*stat*_(*x*|*r*) at *r* < 15 units (specifically, *p*=0.121 for the rejection of the hypothesis of Gaussianity for the subset with *r* between 15 and 16, and *p*>0.25 for subsets with larger *r*, while for the subsets with *r* between 14 and 15 or smaller *r, p*<0.001). This shift of distributions relative to *x*_0_ is biologically meaningful: when a predator is close, depolarization is more likely to occur than hyperpolarization, while when there are no predators nearby, small random fluctuations of the membrane potential *x* are possible in both directions relative to its resting value *x*_0_.

Taken together, the existence and numerical reproducibility of *P*_*stat*_(*r*) and *P*_*stat*_(*x*|*r*) across independent runs of simulations implies the existence of the original stationary distribution *P*_*stat*_(*x,r*), as suggested in Methods. Below, we will use the presented numerical results to introduce an analytical approximation for *u*(*x,r*), valuable for the further analysis.

### Exact solution for the large distance limiting case

We start the analysis of *u*(*x,r*) by considering the limit of *r* → +∞, in which the terms for the sensory perception and the small-distance effective force of the nearest predator distribution vanish, leaving us with only linear (in *x* or *r*) terms in *f*_1_ and *f*_2_:

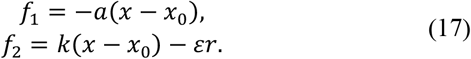

Then, equation (7), taking into account (9), can be rewritten as a differential equation for *u*(*x,r*):

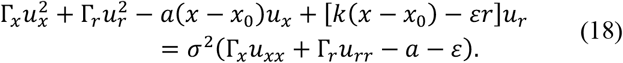

Assuming that the solution is a two-dimensional harmonic potential:

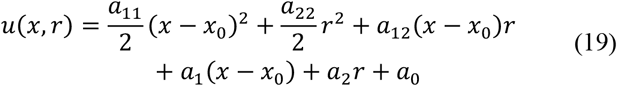

and plugging this form into equation (18), we obtain relationships between coefficients *a*_11_, *a*_22_, *a*_12_, *a*_1_ and *a*_2_ and the parameters of *a, k*, ε, Γ_*x*_ and Γ_*r*_. The solution is given by:

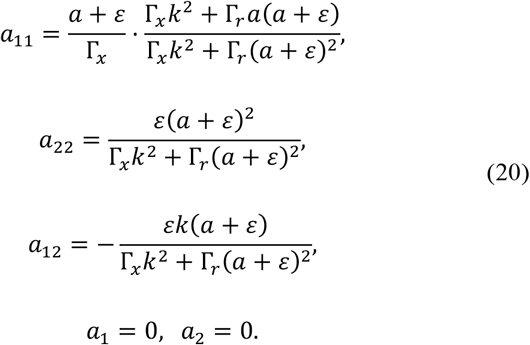

For further derivations, it is convenient to rewrite equation (19) in the following form:

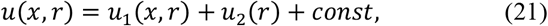

where

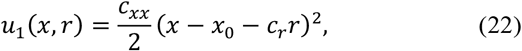

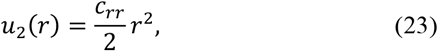

and the coefficients are

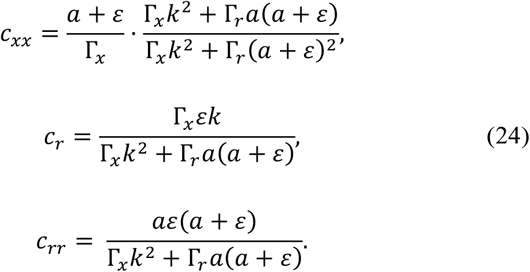

For the parameters used in simulations, as given in equation (14),

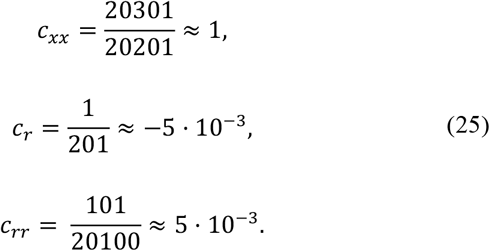

Assuming that *Q* is constant,

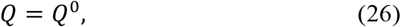

we can find it from the second of equations (11), leading to:

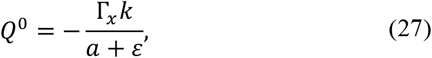

which confirms the assumption on the constant *Q*. Numerically, with the parameters given by equation (14), *Q* equals

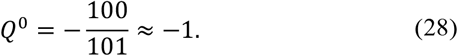

It is straightforward to check that the potential given by equation (21), with the coefficients given by equation (24) and *Q* given by equation (27), correctly predicts the forces given by equation (17).

Note that the exact solution presented here is different from what one might intuitively expect in a mean-field approximation (see Appendix B), emphasizing the importance of strict mathematical analysis, as opposed to verbal reasoning alone. We attract the reader’s attention to this result because the mean-field approximation has been used in this field,^37,41,42,44^ though in ways different than equation (54), sometimes without explicitly checking its accuracy.

### Potential at small and moderate distances r can be approximated with limited changes to the long-distance asymptote

From the analysis of the long-distance behavior of the potential, we proceed to a general case of an arbitrary distance *r*. In this case, we do not have an exact analytical solution. However, all the key conclusions can be made with the use of a certain approximation for the potential *u*(*x,r*) that we will introduce based on the results of numerical simulations reported above.

We assume that the potential *u*(*x,r*) still can be split into two components *u*_1_(*x,r*) and *u*_2_(*r*), as suggested by equation (21), with the terms *u*_1_(*x,r*) and *u*_2_(*r*) resembling those given by equations (22) and (23). In this case, the conditional probability *P*_*stat*_(*x*|*r*) transforms to

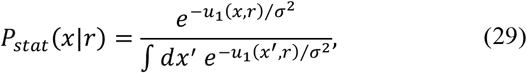

and therefore, for given *r*, the component *u*_1_(*x,r*) as a function of *x* can be found, up to an additive term independent of *x*, as −*σ*^2^ ln *P*_*stat*_ (*x*|*r*). We studied plots of these values as functions of *x* for different narrow intervals of *r* (representative plots are shown in Fig. 3b), and have concluded that a reasonable approximation for *u*_1_(*x,r*) would be:

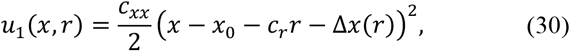

which differs from the long-distance asymptote (22) in that it accounts for a shift of the minimum of the curve for −*σ*^2^ ln *P*_*stat*_ (*x*|*r*) away from the long-distance solution by a value Δ*x* that depends on *r*.

A comparison of the numerical results to the suggested approximation is given in Fig. 3b. Four representative conditional distributions, for the ranges of *r* labeled in green, are shown in four panels. The numerically estimated (from run 1) conditional distribution (on the negative log scale) is shown in red (dots: the most probable values; vertical bars: errors corresponding to the symmetric 5% confidence level, estimated by bootstrapping). The theoretical long-distance asymptote (22) is shown by grey dotted curves, while the approximate formula with the Δ*x* correction (30) is shown by grey solid curves. The resting potential *x*_0_, the minimum of each approximate curve (30) (‘min’) and the Δ*x* correction are also labeled in grey.

At small *r* (Fig. 3b, left top panel), it is clear that the numerical solution corresponds to a curve with a single minimum, though the confidence intervals are wide due to poor statistics. With a bit higher value of *r* (Fig. 3b, right top panel), the sampling improves, and the shape of the potential curve becomes more pronounced, though still somewhat uncertain at large deviations from *x*_0_. The numerically estimated potential curves are noticeably different from the approximate formula in that they are asymmetric and correspond to a broader distribution of *x* than predicted. On the other hand, approximation (30) captures the key properties of the curves, namely, their single-minimum shape and the shift of the minimum by the value of Δ*x*. As we will see below, this degree of approximation is sufficient to draw the key conclusions from this model. At moderate and large *r* (Fig. 3b, bottom panels), the potential becomes harmonic and symmetric in *x* (relative to *x*_0_ – *c*_*r*_*r*). The difference between the numerically generated curve and formula (30) lies within the estimated errors of the former. The difference between the minimum of a potential and the resting potential *x*_0_ becomes negligible at moderate *r* (Fig. 3b, left bottom panel), but re-appears again at large *r* (Fig. 3b, right bottom panel), as expected from equations (22) and (30).

The approximation given by equation (30) is convenient for the analysis, among other reasons, because the integral 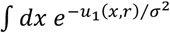 is independent of *r*. Due to this fact, the aggregated probability *P*_*stat*_(*r*) can be written as

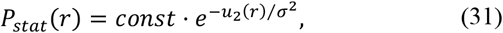

and therefore, the component *u*_2_(*r*) as a function of *r* can be found, up to an additive constant, as −*σ*^2^ ln *P*_*stat*_ (*r*). This component of the potential has a wide minimum, increases gradually at large *r* and sharply at small *r* (Fig. 3c). An accurate fit to the numerical data can be achieved with the following expression (Fig. 3c):

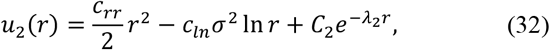

where the first term (quadratic in *r*) is inherited from the asymptote (23), the second term (logarithmic in *r*) is included by analogy with the effective potential (49), where it corresponds to the 1/*r* term in *f*_2_ omitted in the long-distance asymptotic analysis [see equation (17)], and the third term (exponential in *r*) is included based on the numerical analysis (data fit with power law or hyperbolic terms, instead of the exponential term, yielded significantly lower accuracy). This approximation (32) provides high accuracy in a wide range of *r* values (Fig. 3c), including poorly sampled regions of large *r* (due to the use of the exact asymptote) and small *r* (because an exponential function happened to provide an accurate fit for the numerical data). The values of the fitting parameters ensuring the best fit [in the sense of minimal mean squared deviation of ln *P*_*stat*_(*r*)] for the simulations with the chosen parameters (14) were found to be:

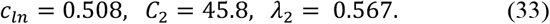

Note that the third (exponential) term in *u*_2_(*r*) plays a critically important role in describing the short-distance behavior of the stationary distribution (at *r* < 10, and particularly at *r* < 5). Without this term, *P*_*stat*_(*r*) would be overestimated by many orders of magnitude at short distances *r* < 5 (Fig. 3c, dotted grey curve).

To complete the construction of an analytical approximation of the potential, consider the dependence of Δ*x* on *r* (Fig. 3c, inset, red dots). In a wide range of *r* values (2 < *r* < 11), Δ*x* exponentially decays with *r* (note the log scale on the Δ*x* axis in the inset). At greater *r*, the numerical estimates of Δ*x* become less reliable due to their small values in comparison to the other component of the shift of the minima, namely, the *c*_*r*_*r* term. A numerical fit of the results of simulations with an exponential function (for the subset of the data with *r* in the range from 2.5 to 10.5) results in:

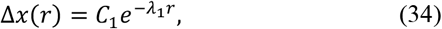

where the best fit for the simulations with the chosen parameters (14) is provided by:

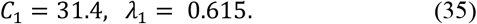

Therefore, the final expression for the approximation of the potential, taking into account equations (21), (30), (32) and (34), assumes the following form:

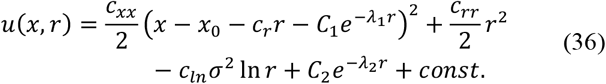

This expression is based on the structure of the long-distance exact result, as given by equations (21)-(23), but takes into account the observations from the numerical simulations (shift of the minimum of −*σ*^2^ ln *P*_*stat*_ (*x*|*r*) as a function of *x* not only at large *r*, but also at small *r*; a term with ln *r* and a negative prefactor, as in the effective potential for the nearest predator; dramatic decrease of probability distribution for *r*, regardless of *x*, at small *r*, as seen from numerical data). On the other hand, this approximation ignores widening and asymmetry of −*σ*^2^ ln *P*_*stat*_ (*x*|*r*) as a function of *x* at small *r*. To check whether this approximation is justified, we further explore what distortions it introduces to the dynamics of the system, namely the forces *f*_1_ and *f*_2_, and whether these distortions could be accepted from the viewpoint of the biological interpretation of the model.

### Suggested analytical approximation of the potential is biologically plausible, given the proper choice of Q

In Appendix C, we derive the function *A* (and therefore *Q*) such that, with the potential given by approximation (36) and taking into account equations (8), the corresponding *f*_2_ will be exact. Then, we use this function *A* to derive the corresponding approximate result for *f*_1_.

A numerical analysis of the resulting *f*_1_ demonstrates a biologically acceptable behavior at biologically meaningful values of *r* and *x*. The approximate solutions for different *x* go side-by-side, plateau at large *r*, and rapidly increase at small *r*, exactly like the exact desired expressions for *f*_1_ (Fig. 4a). Further analysis can be based on the split of the resulting expression for *f*_1_ into three parts (Appendix C):

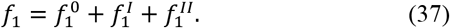

**Fig. 4.**
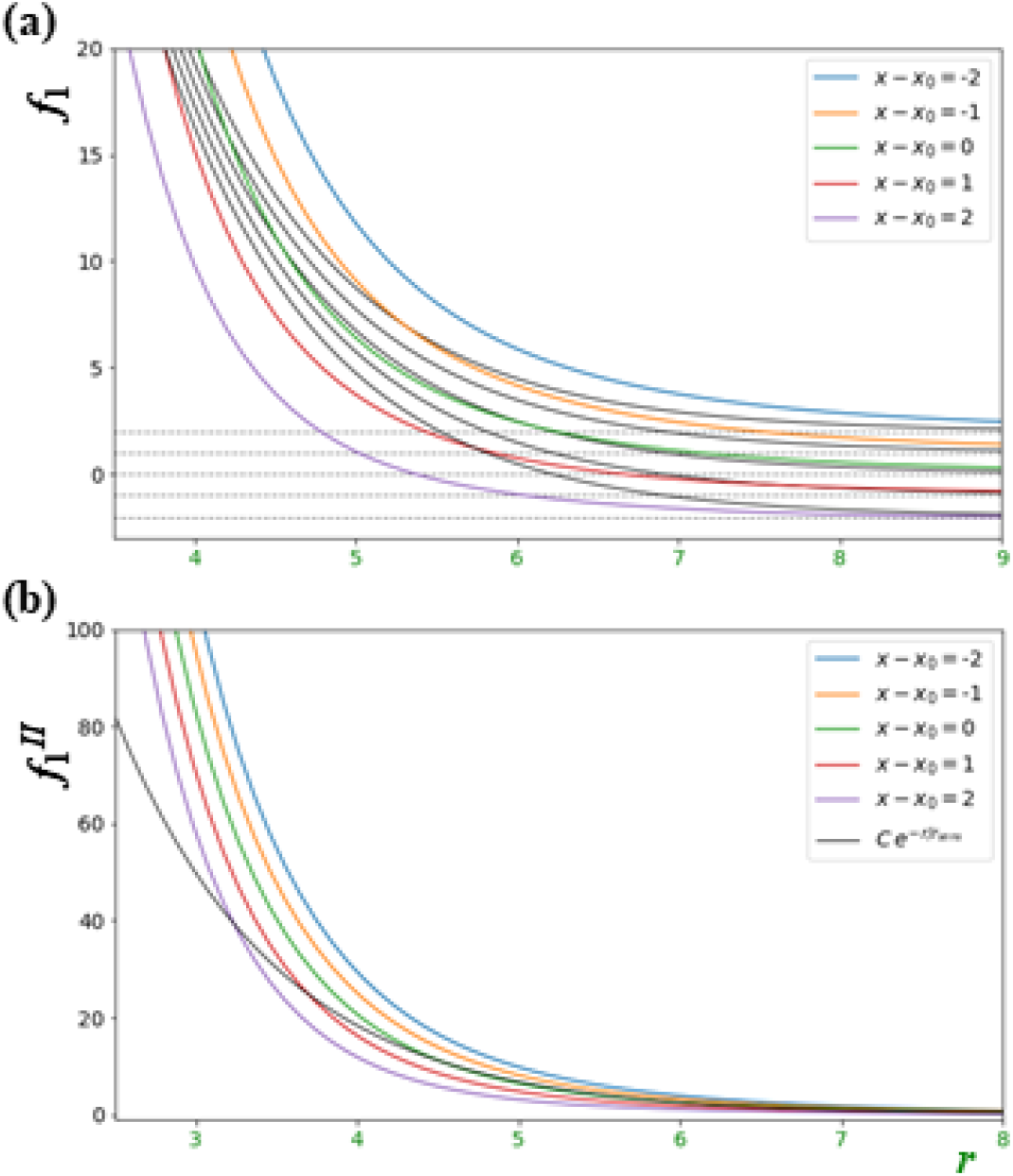
Approximation for the potential *u*, equation (36), given the proper choice of *A*, is biologically plausible. (a) Approximate *f*_1_ (*multicolored curves*) computed from equation (70) assuming 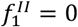 behave qualitatively the same as the exact *f*_1_ defined by equation (2) (*nearest black curves*) and possess the same large-distance asymptotes (17) (*dotted grey curves*). Specifically, at large distances, *f*_1_ plateau at an *x*-dependent level (biological interpretation: when predators are at a large distance away, a neuron gets no sensory input, while sensory-independent mechanisms return the membrane potential to its resting value); at shorter distances, *f*_1_ increase (a neuron starts feeling an approaching predator, and the membrane potential increases); at even shorter distances, *f*_1_ rapidly increase (the closer the predator, the greater the sensory input). Both the exact and approximate curves go in bundles side-by-side, in the order of increasing *x*, and without crossings (given a constant distance to the predator *r*, the rate of change of the potential *x* monotonically decreases with *x*). (b) The contribution 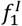 to *f*_1_ slightly depends on *x* (*multicolored*), while the exact sensory term 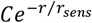 is the same for all *x* (*black*). Also, at *r* = 0 the exact term is finite, while the approximate 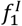 diverge to infinity. However, in the range of sampled values of *r* (Fig. 3a, *green*), both differences are biologically immaterial. Note that the other contribution to *f*_1_, namely 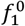, coincides with the exact one (see the text).

The zeroth term 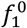 coincides with the long-distance result (17):

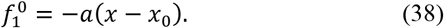

The biological interpretation of this term is that there are certain mechanisms (such as opening of certain voltage-gated ion channels) that return the membrane potential *x* to its resting value *x*_0_, and that these mechanisms work in the same way regardless of the distance to the predator *r*.

Having written out explicit expressions for 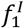, one can check that it vanishes in the limit of *r* → +∞, as desired. At finite *r*, the comparison of the approximate solution with *u* and *A* given by equations (36) and (63), respectively, and the exact desired sensory term of 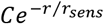, as suggested by equation (2), shows the following (Fig. 4b). The contribution 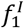 not only decays in the limit of *r* → +∞ in principle, but becomes practically negligible at moderate values of *r* ∼ *r*_*typ*_. Also, it rapidly grows at small *r*, like the desired exponential term, being comparable to 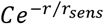 at sampled short distances *r* ∼ 3-5. Two noticeable differences between 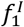 and 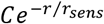 are: (1) a faster growth of 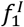 with decreasing *r* at *r* < 3 (in particular, in the limit of *r* → 0, the exponential term reaches a large, but finite value *C*, while 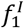 diverges to +∞), and (2) a dependence of 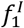 on *x* (Fig. 4b). However, these differences are insignificant from the viewpoint of the biological interpretation of the model. The region of *r* values of *r* < 3 is rarely sampled (Fig. 3a), and for the plausible behavior of the model, it is sufficient that 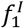 rapidly increases as *r* decreases in this range of *r* values, while the quantitative accuracy is not vital. As for the dependence of 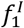 on *x*, in the range of biologically plausible values of *x*, this dependence does not interfere with the sensory character of the neuron, and only modulates it. A somewhat higher sensitivity of the neuron to an external signal (predator approach) if this neuron is already in an excited state, *x* > *x*_0_, and a somewhat lower sensitivity if it is in an inhibited state, *x* < *x*_0_, resembles sensitization, a well-known phenomenon in neurons. Though we originally did not intend to include this phenomenon into the model, its appearance in the approximate solution does not look implausible and does not seem to affect the major conclusions we draw in this work.

We chose this direction of reconstruction of *f*_*i*_ – from *f*_2_ to *f*_1_ – because an appearance of a weak dependence of one of the four terms 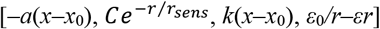 on the other variable is more acceptable for the sensory term 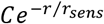, as explained above. Both terms in *f*_2_ do not allow such an admixture of the other variable. The term *k*(*x*–*x*_0_) stems from the activation of effectors by the nervous system, and it would be more difficult to imagine a molecular mechanism that would cause this motor response to be affected by the distance to the predator. The term *ε*_0_*/r*–*εr* is set by the properties of the environment (spatial distribution of predators) and cannot be directly changed by the nervous system excitation. Therefore, it is more reasonable to build the approximation in such a way that the *f*_2_ terms are predicted exactly, while the approximation error is localized to the sensory term.

The remaining term 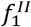 is discussed in the next subsection.

### Small sigma approximation may lead to qualitatively incorrect results

In the literature, it is common to omit the terms with partial derivatives of *Q*, justifying it by a small *σ*^2^ (weak noise) expansion.^18, 27-29, 35, 36, 40, 43-45^ Here, we draw the reader’s attention to three issues in the presented simple model that this approximation causes.

First, this approximation yields a qualitatively incorrect result for the reconstruction of *f*_1_ from *f*_2_, given the potential *u*, unlike the successful reconstruction presented in the previous subsection. Accepting approximation (12), *Q* can be expressed from *f*_2_ and *u* in the following way:

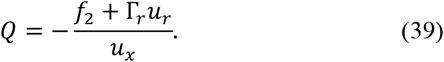

Plugging this expression into the equation for *f*_1_, we get:

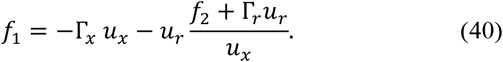

For the approximate potential (36), successfully used in the previous subsection to reconstruct a biologically plausible *f*_1_ (Fig. 4), the partial derivative *u*_*x*_ equals zero when 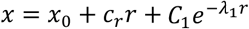, that is, at the value of *x* at which *u*(*x,r*) reaches a minimum as a function of *x* for every *r*. In other words, the denominator in the right-hand side of equation (40) is nullified at the most probable value of *x* at each value of *r* (including all highly probable ones). A straightforward check shows that the numerator at such values of *x* and *r* is non-zero. Thus, the function *f*_1_ derived under the assumption of vanishing “small *σ*^2^ terms” – unlike one derived under the exact treatment – is quantitatively wrong, diverging to infinity at the most probable values of its arguments.

Second, with approximation (12), we concluded that *Q* (and therefore *A*) is uniquely determined by *f*_2_, as shown in equation (39). However, the exact treatment in the previous subsection demonstrated that *A* (and hence *Q*) is defined, given *f*_2_, up to an integration ‘constant’ *C*(*r*) [see equations (63) and (67)]. Consequently, *f*_1_ is defined up to an additive term 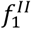 [see equations (37) and (73)]. With the potential given by equation (36), 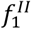 can be written as

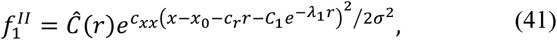

where *Ĉ*(*r*) is another arbitrary function of *r* defined by 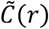 and *u*_2_. In the limits of *x* both much above or much below *x*_0_, this component 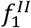 has the same sign (whether it is positive or negative, depends on *Ĉ*(*r*)), introducing a bias toward higher or lower values of *x*. This term, however, is finite for *x* in the plausible range of its values, and by construction should not change the stationary distribution, which is controlled by the potential *u*(*x,r*). Though this component 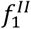 has a somewhat unclear biological interpretation – and for this reason we omitted it in the previous subsection – we nevertheless think that the existence of this contribution, to a large degree arbitrary, should be acknowledged on the conceptual level.

Third, note that the limit of *σ*^2^ → 0 implies, according to equation (9), that *P*_*stat*_(*x,r*) turns into a Dirac delta function *δ*(·) centered on the global minimum of the potential function *u*:

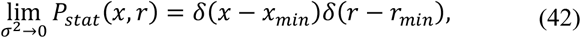

where

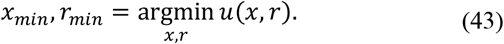

In particular, for the approximate potential (36),

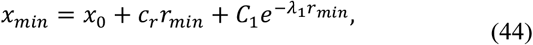

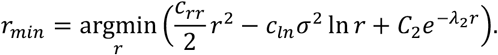

Then, the behavior of the potential *u*(*x,r*) at all values of *x* and *r* different form the global minimum turns out to be irrelevant and does not affect the predictions of the model. Therefore, in the limit of *σ*^2^ → 0, we cannot make any exact conclusions about the states of the neural network or the environment, except for the only state with the highest probability density at steady state (and perhaps its infinitely small vicinity).

Thus, neglecting terms of the *σ*^2^ order (and higher orders) may lead to qualitatively incorrect results in the described theoretical framework at least in three possible aspects listed above.

## Discussion

In this study, we have presented a highly simplified (perhaps the simplest) model of the nervous system. Our primary intention behind its design was to facilitate its role as a template, accommodating both mathematical derivations of relevant equations and numerical modeling.

Our inspiration for this model stemmed from a hypothetical scenario concerning the origin of the nervous system.^2^ We hope that in upcoming research, this avenue of exploration may provide insights into the genesis of the nervous system. However, our current rendition of the model should not be regarded as an accurate representation of the primordial nervous system. Rather, our primary focus was to ensure that our model encompasses all pivotal facets proposed by the hypothetical scenario. It is our earnest hope that substantial segments of our presented model transcend the confinements of this hypothetical situation. This is especially pertinent to the formalism we employed, which revolves around using the potential *u* and the function *A* (or *Q*) as the independent controls defining the dynamics of a nervous system within a stipulated environment.

Crafting a rudimentary yet holistic model presents inherent challenges. Foremost among these challenges is determining which features warrant retention, versus those which can be sidelined. Our model takes into account both neuronal and environmental dynamics (retaining one degree of freedom for each) along with the governing laws of each dimension. We also incorporated evolutionary aspects, namely the predator evasion. Contrarily, we excluded elements such as a multi-neuronal network, specialization of neurons (e.g., to sensors, effectors, interneurons), the intricate dynamics of spiking neurons that demands more degrees of freedom and the details on the molecular mechanisms supporting the introduced dynamics of the system. Whether these choices are justified should be decided, in our opinion, based on the capability of the model to inspire novel research, rather than their formal consistency.

One of the strengths of our model lies, in our opinion, in its simplicity and robustness, both from numerical and analytical standpoints. Theoretically, we can confidently expound on certain facets using mathematical formulations. These encompass the existence of the stationary distribution, the potential and the associated stationary Fokker-Planck equation, along with an explicit expression for drift terms *f*_*i*_ in terms of *u* and *Q*. Our numerical simulations affirm that the model allows for exhaustive sampling even with limited computational resources. Guided by these simulation results, we were able to derive approximate formulas corresponding to biologically plausible solutions. Collectively, our model, coupled with its exact, numerical and approximate solutions, has the potential to be an instrumental tool in modeling neural networks.^46,47^ He hope that its importance for future research might be equated with simple models like the hydrogen atom model in quantum chemistry or the Lorenz system in dynamical systems theory.

Our explorations with this model yielded some intriguing findings. Notably, we identified significant constraints of the small-sigma limit, including that it may result in biologically implausible dynamics, with *f*_1_ diverging infinitely at most probable states of the system. Also, the exact solution derived for the effective potential differs from intuitive expectations, which underscores the importance of rigorous mathematical treatment over mere heuristic reasoning. Such insights, though manifestly evident within our model, have not been paralleled or deduced in past research, to the best of our knowledge. The insights gleaned from this study lay the groundwork for upcoming investigations, potentially focusing on facets like evolutionary optimization.^46^

In conclusion, we present a simple model that, despite its inherent abstract nature, may serve as a promising framework to explore critical facets of neural networks that reveal themselves even in the simplest nervous systems. As with any model, there are limitations, but the strengths seen here lay the foundation for further research, especially as articulated in the companion paper^32^ discussing evolutionary optimization. Our findings also serve as a cautionary tale against the over-reliance on simplistic approximations, such as the weak-noise or mean-field approximations, emphasizing the need for in-depth mathematical analyses to capture the true essence of neurobiological dynamics.

## Appendices

### Appendix A. Effective field for the nearest predator distribution

Assume that other “Dickinsonia” (“predators”) are evenly distributed in space and do not actively pursue the victim (in other words, are predators only by accident). It is reasonable to consider their mutual motion in an effectively two-dimensional space, because they approach toward and move away from each other along two horizontal dimensions, while the vertical dimension serves only to switch between the feeding and floating regimes.

Denote the average density of predators, in pieces per unit area, as *ρ*. Then a disc around the organism with the radius of *r* has an area of *πr*^2^ and, therefore, contains on average *πρr*^2^ predators. The probability that in fact it contains exactly *N* predators is described by the Poisson distribution:

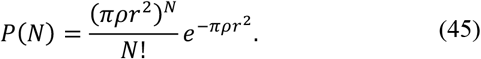

The probability that this disc does not contain any predators can be obtained by letting *N* = 0:

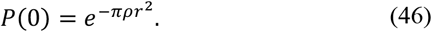

Therefore, the probability that the closest predator is at the distance not exceeding *r* is equal to

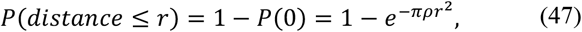

which is a cumulative probability. The probability density can be found by differentiating this cumulative probability:

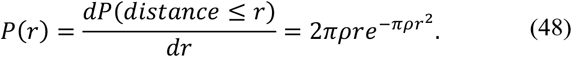

Then, by analogy with equation (9), the effective potential that corresponds to this distribution is

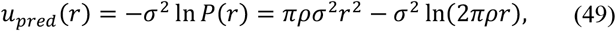

and, by analogy with equation (11), the corresponding contribution to *f*_2_ is

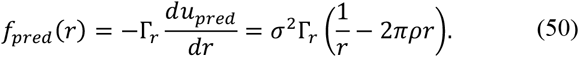

Finally, it is convenient to rewrite this equation in terms of the distance *r*_*typ*_ at which the probability density is maximal or, equivalently, *f*_*pred*_ vanishes:

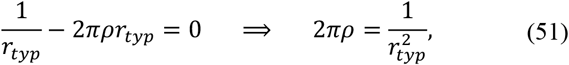

which leads to:

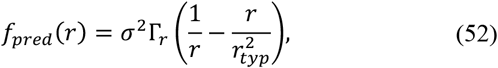

or, even simpler, as

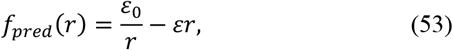

where *ε*_0_ = *σ*^2^Γ_*r*_ and 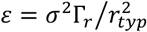 are constants.

### Appendix B. Naïve estimate of the long-distance behavior of the potential and Q

To emphasize the importance of strict mathematical treatment, we contrast the exact results for the large-distance limit, equations (19)-(24), to a plausible intuitive guess about that solution that one might come to (at least, the equations below were our first guess, before we derived the exact solution).

At large distances, we could expect that the neuron did not sense the predator and did not initiate an escape response, and hence, the distributions of membrane potential and distance to the predator were independent of each other (a mean-field approximation):

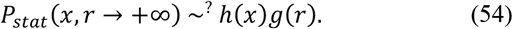

To be compatible with the derivation given in Appendix A, the *r*-dependent component should satisfy the following equation:

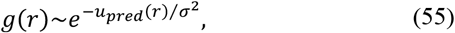

where *u*_*pred*_ is the effective potential given by equation (49). In a combination with

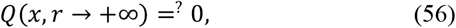

which could be justified by reasoning that a system with two independent degrees of freedom was not dissipative, this reproduced the exact desired result on the *f*_2_ term:

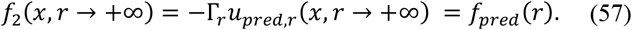

However, the naïve guess for the potential given by equation (54) contradicts to the exact results given by equations (21)-(24). In particular, the leading term in the potential, corresponding to equations (54) and (55), at large *r* is *εr*^2^/2, while the exact result has the leading term of *c*_*rr*_*r*^2^/2 (Fig. 3c). These prefactors differ by *ε*/*c*_*rr*_, which for the parameters used in the simulations [see equations (14) and (25)] equals 201/101 ≈ 2. Also, the true value of *Q*, as given by equations (26)-(28), is different from zero. Numerical results discussed in the main text confirm the long-distance asymptote (21)-(24), not the naïve estimate presented here. Therefore, an intuitive guess about the solution asymptote at large distances proves to be incorrect, underscoring the importance of strict mathematical analysis, as opposed to solely verbal reasoning.

### Appendix C. Solution for the function A(x,r) providing the exact result for f_2_ with the approximate potential u(x,r)

Let us find the function *A* (and therefore *Q*) such that, with the potential given by approximation (36), the predicted *f*_2_ will be exact. As follows from equation (8), this requirement leads to:

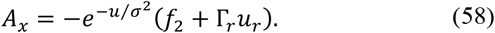

Integrating over *x*, we obtain:

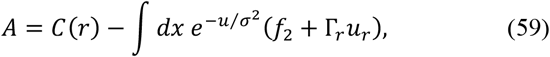

where *C*(*r*) is an integration constant, but it appears from the integration over *x*, so it may depend on the other independent variable *r*. Since only the first term in equation (36) depends on *x*, the exponential of all other terms in *u*(*x,r*) can be moved out of the integral, resulting in:

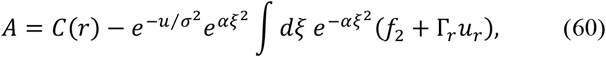

where

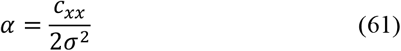

and

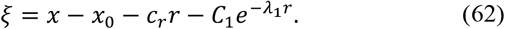

Using the exact expression for *f*_2_ given by equation (3) and the partial derivative *u*_*r*_ computed from the potential (36), both rewritten in terms of *ξ* (62), we can calculate the integral over *ξ* analytically:

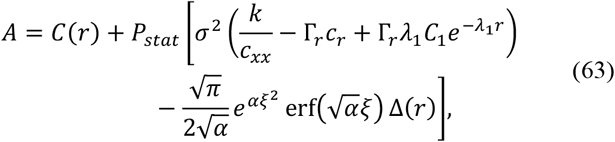

where

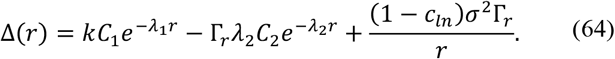

Then, *f*_1_ can be expressed from equation (8) as

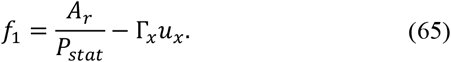

For the convenience of further analysis, we split *A* into three terms:

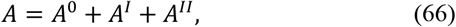

where we arrange the terms in equation (63) into these three components of *A* in the following way:

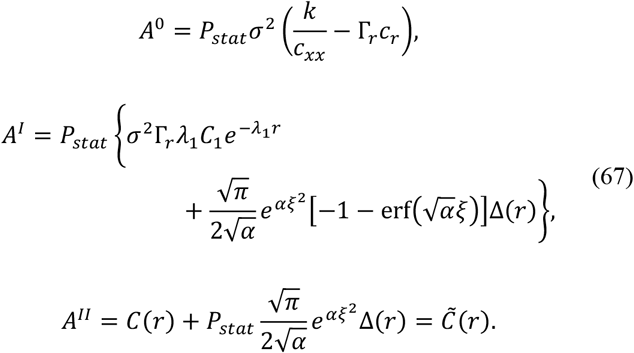

The zeroth term *A*^0^ corresponds, according to equation (9), to a constant contribution to *Q*, namely *Q*^0^ = − *A*^0^/*σ*^2^*P*_*stat*_ = Γ_*r*_*c*_*r*_ − *k*/*c*_*xx*_. Plugging in the expressions for *c*_*r*_ and *c*_*xx*_ from equations (24), we can simplify this to *Q*^0^ = − *k*Γ_*x*_/(*a* + *ε*), which coincides with equation (27) derived in the long-distance asymptotic analysis. Therefore, the term *A*^0^ in equation (66) corresponds to the long-distance asymptotic solution for *A*.

The term *A*^*I*^ provides a correction to *Q* from the viewpoint of large *r* series expansion. We have introduced “–1” before the error function in it to ensure a convenient long-distance asymptotic behavior (the corresponding counter-term was added to *A*^*II*^). Specifically, at *r* → +∞ and finite *x, ξ* → −∞, and therefore

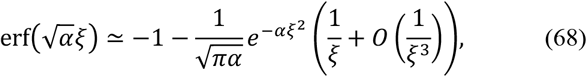

which yields the following long-distance asymptotes for *A*^*I*^:

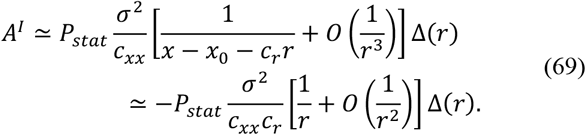

Hence, the term *A*^*I*^ provides a correction to the zeroth term *A*^0^ decaying at long distances *r* by a power law.

As for the last term *A*^*II*^, the product 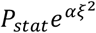, as noted above, depends only on *r*, but not *x* or *ξ*, so the whole expression for *A*^*II*^ also depends only on *r*. Therefore, *A*^*II*^ can be interpreted just as a redefined integration constant, which we emphasize by the notation 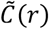.

Plugging the expressions for *A* and its components, equations (66)-(67), into the equation (65), we obtain, after some transformations, the following result for *f*_1_:

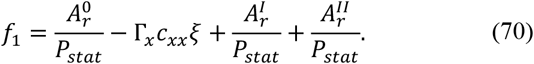

Next, we split this expression for *f*_1_ into three components, as suggested by equation (37). We define the zeroth component as those terms in equation (70) that emerge from *Q*^0^ and Γ_*x*_ *u*_*x*_ and do not vanish in the long-distance limit:

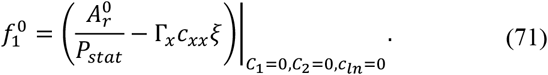

After simple transformations, taking into account equations (24), we conclude that the expression for 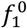 simplifies to 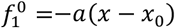, which coincides with the long-distance asymptote (17).

We define the second component as the difference between the exact expression for the first three terms in equation (70) and the expression for 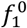:

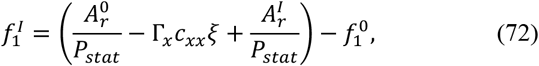

while the last contribution to *f*_1_ is defined directly in terms of the corresponding contribution to *A*:

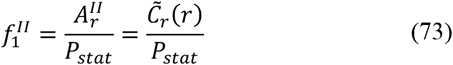

(we did not follow this pattern for the first two terms to maintain the connection between the zeroth order terms and the exact expressions from the long-distance analysis, and hence 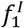 includes contributions not only from *A*^*I*^, but also from *A*^0^ and *ξ*).

To finish the discussion of the derivation of *f*_1_, we note that if

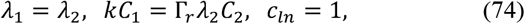

then Δ(*r*) vanishes, and the expression for *A*^*I*^ (67) simplifies (in particular, the corresponding *Q*^*I*^ reduces to a single term exponentially decaying with *r*). Respectively, the analytical expression for *f*_1_ also simplifies, making its analysis easier. However, the behavior of the resulting *f*_1_ is quantitatively the same as shown in Fig. 4: this simplification neither creates biologically irrelevant artifacts, nor eliminates the two above-mentioned differences between the approximate and the originally intended solutions. Numerically, comparing conditions (74) to the empirically found values (33) and (35), we conclude that the first two conditions in (74) are approximately satisfied, while the third condition, on *c*_*ln*_, is far from being justified by the numerical data. We found that enforcing the condition of *c*_*ln*_ = 1 during the numerical fit of *u*_2_(*r*), equation (32), to the results of simulations (Fig. 3c) significantly deteriorates the quality of the fit. With a value of *c*_*ln*_ different from 1, Δ(*r*) retains the 1/*r* term, and 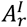 remains complicated. For these reasons, we did not enforce conditions (74) in the presented analysis.

